# High dimensionality of the stability of a marine benthic ecosystem

**DOI:** 10.1101/2020.10.21.349035

**Authors:** Nelson Valdivia, Moisés A. Aguilera, Bernardo R. Broitman

## Abstract

Stability is a central property of complex systems and encompasses multiple dimensions such as resistance, resilience, recovery, and invariability. How these dimensions correlate among them is focus of recent ecological research, but empirical evidence at regional scales, at which conservation decisions are usually made, remains absent. Using a field-based manipulative experiment conducted in two marine intertidal regions, we analyse the correlations among different aspects of stability in functioning (community cover) and composition of local communities facing a press disturbance. The experiment involved the removal of the local space-dominant species for 35 months in eight sites under different environmental regimes in northern- and southern-central Chile (ca. 30°S and 40°S, respectively). After the disturbance, the magnitude of the initial responses and the recovery patterns were similar among communities dominated by different species, but varied between the functional and compositional response variables, and among four dimensions of stability. The recovery trajectories in function and composition remained mostly uncorrelated across the system. Yet, larger initial functional responses were associated with faster recovery trajectories—high functional resilience, in turn, was associated with both, high and low variability in the pattern of recovery. Finally, the compositional stability dimensions were independent from each other. The results suggest that varying community compositions can perform similar levels of functioning, which might be the result of strong compensatory dynamics among species competing for space in these communities. Knowledge of several, and sometimes independent, aspects of stability is mandatory to fully describe the stability of complex ecological systems.

## Introduction

Anthropogenic environmental change is disrupting the functioning and composition of ecosystems over multiple spatiotemporal scales (De Laender et al., 2016; Isbell et al., 2017). Understanding the capacity of ecosystems to absorb and respond to such disturbances is challenging, because stability is a multifaceted property characterised by several dimensions (Pimm, 1984; Ives and Carpenter, 2007; Donohue et al., 2013). Recent theoretical and empirical advances provide the framework to assess this dimensionality through correlations between multiple aspects of stability such as resistance, resilience, recovery, and invariability (Donohue et al., 2013; Hillebrand et al., 2018; Radchuk et al., 2019). Most scientific research to date has focused on either one or two dimensions, which has impaired our capacity to fully understand the stability of complex ecological systems (Donohue et al., 2016; Kéfi et al., 2019). In addition, the functional consequences of biodiversity loss can be exacerbated at regional spatial scales, which are relevant for conservation or policy (Isbell et al., 2017). To our best knowledge, however, the correlations of multiples dimensions of stability have not been assessed at large spatial scales yet (Gonzalez et al., 2020).

Each dimension of stability can be assessed either for aggregate ecosystem properties performed by the whole community, such as productivity, community biomass, or cover (functional stability), or for the combination of species identities and abundances in the community (compositional stability; Micheli et al., 1999). The relationship between both domains of stability sheds light into the contribution of particular combinations of species to the aggregate function. For example, if the community is characterised by asynchronous species dynamics, in which the decay of some species is compensated by proportional increases of others, then the community could maintain relatively stable aggregate properties over time (Yachi and Loreau, 1999; Gonzalez and Loreau, 2009). This implies that a low compositional stability can sustain a high functional stability (Tilman, 1996; Micheli et al., 1999). In these cases, negative or null correlations between functional and compositional aspects of stability can be observed (e.g., Hillebrand et al., 2018; Hillebrand and Kunze, 2020). The stabilising role of such asynchronous or compensatory species dynamics is supported by several studies (Gonzalez and Loreau, 2009; Valencia et al., 2020). However, it is less clear how functional and compositional stabilities will be correlated in communities facing the widespread chronic disturbances experienced by ecosystems (Guelzow et al., 2017).

Within the functional and compositional domains, the dimensions of stability include, but are not limited to, *resistance*, or the degree to which an aggregate function or species composition remains unchanged after a disturbance, *resilience*, or the speed of community recovery after the disturbance, *recovery*, or the degree to which the community recovers from the disturbance, and *invariability*, or the degree of constancy of the function or composition around the temporal recovery trend (Grimm and Wissel, 1997; Hillebrand et al., 2018). These aspects of stability can be independent of each other or can be correlated. Several models suggest that most stability dimensions are positively correlated (e.g., Scheffer et al., 2009; Radchuk et al., 2019). For instance, communities with low invariability (i.e., highly variable functioning or composition over time) can show low resistance against disturbances, because of the increased risk of extinctions of species with small population sizes (Lande et al., 2003). In those cases, stability is unidimensional and conservation decisions could maximise both aspects of stability, i.e., enhancing resistance would also enhance invariability. However, negative correlations among dimensions can be also observed. Resistance, for example, can be negatively related with resilience when disturbance has a strong impact on the community and intrinsic growth rates are high: Larger initial disturbances can allow for faster recovery trajectories in function or composition (Harrison, 1979; Hillebrand et al., 2018). Negatively correlated stability dimensions imply that policy trade-offs would be needed to maintain either one or another aspect of stability. Finally, chronic disturbances, like the loss of dominant species, can lead to uncorrelated dimensions of stability (Donohue et al., 2013). The lack of correlation among stability dimensions (i.e., a high dimensionality) indicates that multiple aspects need to be quantified simultaneously to fully characterise the overall stability of the ecosystem.

In this study, we analyse the results of a field-based manipulative experiment in which a press disturbance, i.e., the chronic extinction of the locally dominant species, was simulated to assess the dimensionality of stability within the functional and compositional domains of rocky intertidal sessile communities. To this end, we used total community cover as a relevant metric of ecosystem functioning. For communities of sessile species, this function expresses the degree to which competing species use the space as a limiting resource and thus, its relevant to understand both functional and compositional stability (Paine, 1980; Dunstan and Johnson, 2006; Stachowicz et al., 2008). Multiple sites spanning a large environmental gradient (Fig. 1) were used to assess the dimensionality of stability at an ecologically relevant spatial scale (Isbell et al., 2017). Our experimental design involved that the locally dominant species varied among sites, which allowed us to enhance the generality of our findings (e.g., Arnillas and Cadotte, 2019). These dominant species were functionally and evolutionary distinct and included the red corticated alga *Mazzaella laminarioides*, chthamalid barnacles (mixture of *Jehlius cirratus* and *Notochthamalus scabrosus*), and the purple mussel *Perumytilus purpuratus* (hereafter referred to as *Mazzaella*, barnacles, and *Perumytilus*, respectively; Table 1). This setup allowed us to test the following three hypotheses.

**Fig. 1.**
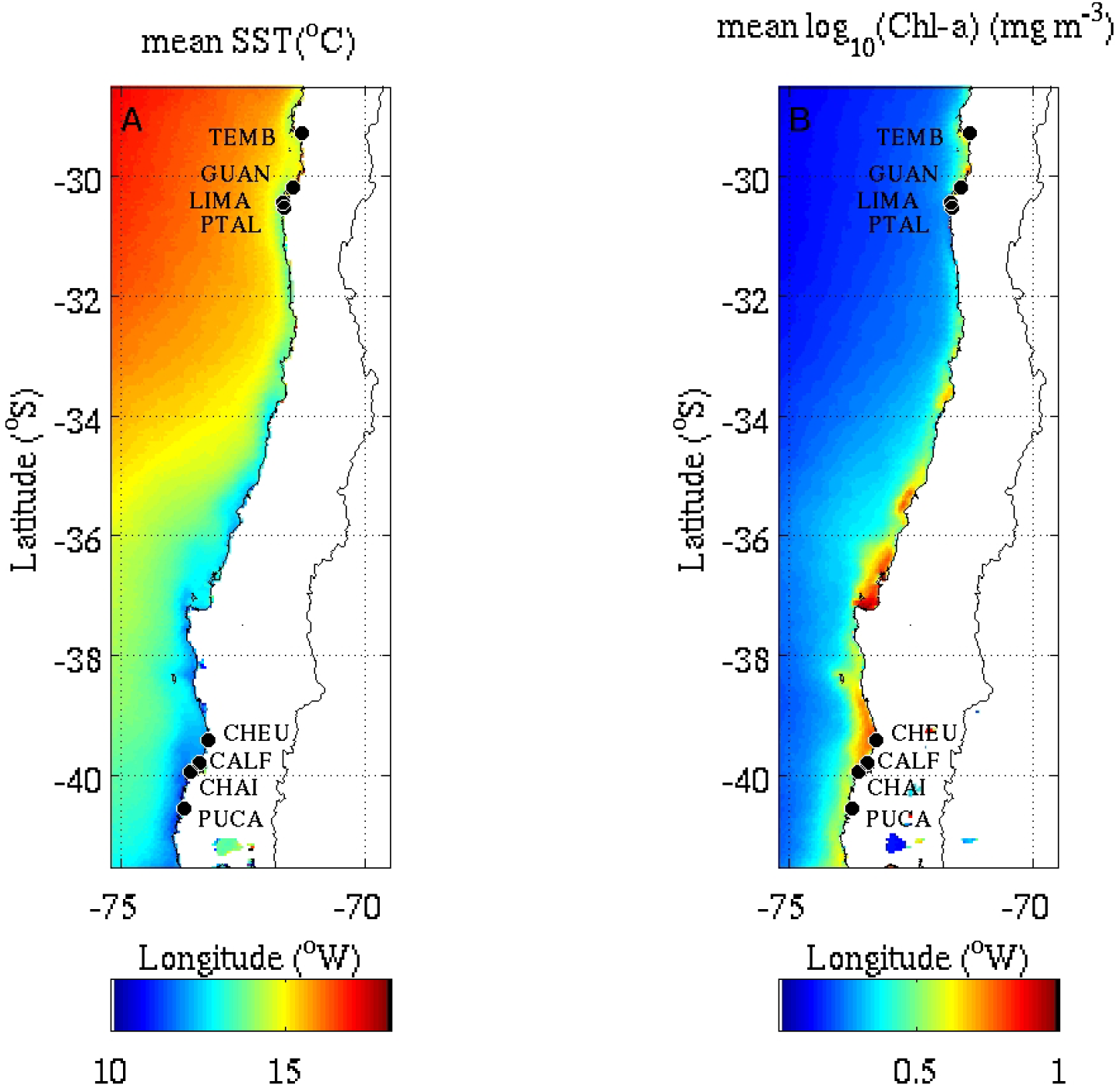
Location of study sites in the northern and southern regions in the SE Pacific shore of Chile. Colour scales denote long-term temporal mean (2003-2018) of (A) sea surface temperature and (B) log_10_ Chlorophyll-*a* concentration using L3 MODIS-aqua data (Lara et al., 2019). Site codes are in Table 1. *Mazzaella*-dominated communities: LIMA and PTAL; barnacle-dominated communities: TEMB, GUAN, and CHAI: *Perumytilus*-dominated communities: CHEU, CALF, and PUCA. Sea surface temperature and Chlorophyll-a data were not included in the statistical analyses and are shown here to describe the environmental conditions across the province.

**Table 1.** Identity of the dominant species experimentally removed at each rocky shore site in each region. South latitude and West longitude (Lat. and Lon. respectively) data are provided in decimal degrees. Community-specific mean percentage (1^st^ and 3^rd^ quantiles parenthesised) cover of each locally space-dominant species (*Mazzaella laminarioides*, mixture of *Jehlius cirratus* and *Notochthamalus scabrosus* barnacles, and *Perumytilus purpuratus*) is provided. Simpson’s concentration index (λ; range: [0, 1]) indicates the degree of unevenness in the distribution of species abundances. Community-specific mean percentage covers are in bold.

| Region | Lat. | Lon. | Site | Site<br>code | *Mazzella* | Barnacles | *Perumytilus* | $\lambda$ |
| --- | --- | --- | --- | --- | --- | --- | --- | --- |
| Northern | 29.28 | 71.18 | Temblador | TEMB | 0 (0-0) | **40** (25.5-56.2) | 0.2 (0-0) | 0.43 |
|  | 30.20 | 71.48 | Guanaqueros | GUAN | 0 (0-0) | **69** (67.5-80.2) | 0.3 (0-0.8) | 0.67 |
|  | 30.45 | 71.42 | Limarí | LIMA | **54.3** (41.2-65) | 0.5 (0-1) | 0 (0-0) | 0.48 |
|  | 30.54 | 71.41 | Punta Talca | PTAL | **31.1** (19.5-48.5) | 1.3 (1-2) | 0.1 (0-0) | 0.37 |
| Southern | 39.41 | 73.22 | Cheuque | CHEU | 2.3 (0-0) | 21.9 (2-40) | **100** (100-100) | 0.59 |
|  | 39.78 | 73.39 | Calfuco | CALF | 7.6 (0-2.2) | 4 (1-5.2) | **100** (100-100) | 0.62 |
|  | 39.93 | 73.59 | Chaihuín | CHAI | 7 (0-8) | **89** (84-97.2) | 3.5 (1-3.5) | 0.77 |
|  | 40.54 | 73.72 | Pucatrihue | PUCA | 0 (0-0) | 1.1 (1-1) | **98.5** (98.8-100) | 0.86 |

### H1: The magnitude of the stability response will depend on the identity of the removed dominant species

The three dominant species use the settlement substratum in different ways: *Perumytilus* is able to monopolise almost the 100 % of primary substratum, while the barnacle complex occurs in combination with bare rock and filamentous algae (Table 1; Broitman et al., 2001; Moreno, 2001; Lagos et al., 2005). *Mazzaella*, on the other hand, uses a smaller proportion of the primary substratum, but generates a large canopy-like structure above the rock (Hoffmann and Santelices, 1997). Thus, the removal of these species should trigger different responses of the ensuing assemblages (see also next paragraph, below).

### H2: Across sites, the removal of the dominant species will elicit negative or neutral correlations between functional and compositional stability dimensions due to prevailing compensatory dynamics

The removal of *Perumytilus* triggers the compensation between subordinate taxa: fast-growing ephemeral macroalgae such as *Ulothrix* sp. and *Pyropia* spp. dominate the assemblage during the first two months after the removal, then they are replaced by barnacles and *Ulva* sp., and eventually by *Mazzaella* (Aguilera and Navarrete, 2012; Tejada-Martinez et al., 2016). Barnacles facilitate small grazers; therefore, the removal of barnacles leads to increased abundances of microalgae but decreased abundances of perennial macroalgae (Harley, 2006; Mrowicki et al., 2014; Whalen et al., 2016). Finally, the removal of *Mazzaella* is associated with the decay of the red alga *Gymnogongrus furcellatus* (*Asterfilopsis furcellata*), but the increase in the abundance of *Ulva rigida, Lithophyllum* spp., and the ephemerals *Petalonia fascia* and *Scytosiphon lomentaria* (Jara and Moreno, 1984)—these dynamics are ascribed to shading effects of *Mazzaella* on the understorey (Moreno and Jaramillo, 1983; Nielsen and Navarrete, 2004).

### H3: Across sites and within the stability domains (functional and compositional), the stability dimensions will be correlated with each other

According to previous modelling work (Radchuk et al., 2019), the stability dimensions will show positive pairwise correlations. The exception will be negative correlations of resistance-resilience, because stronger impacts can allow faster recovery rates, and resilience-invariability, because a faster recovery is expected to be associated with more variable temporal trajectories in function and composition (Hillebrand et al., 2018).

## Materials and methods

### Study province

The study was conducted at multiple field sites on the extremes of a large transitional biogeographic province that extends between the Peruvian and Magellan provinces along the Chilean sector of the South East Pacific (Camus, 2001; Thiel et al., 2007; Lara et al., 2019). We used eight study sites that where evenly split between the northern and southern extremes of the province (around 30.23°S and 39.9°, respectively)—each region encompassing ca. 200 km of shoreline (see Fig. 1). Therefore, the experimental design considered four spatial scales: The province encompassed two regions; each region included four sites. Then, each site contained several replicated plots. A plot was a fixed location where the abundance of macrobenthic sessile species was estimated over time. There were two types of plots, either control or disturbed in which the dominant species was removed. Within each site, each stability metric was calculated for randomly paired plots (one of each type of plot). Depending on the tested hypothesis, the analyses were conducted at the scale of individual plots, paired plots, or after aggregating these data according to the identity of the removed species, which was a site-level attribute.

The northern region is characterised by semi-permanent coastal upwelling activity, which coincides with a significant transition in abiotic (Hormazabal et al., 2004; Garreaud et al., 2011; Tapia et al., 2014) and biotic conditions (Valdivia et al., 2015). Compared with neighbour locations, this upwelling point has been associated with more persistent abiotic environmental conditions, more synchronous species fluctuations, and lower invariability in total cover (Valdivia et al., 2013).

The southern region presents significant correlations between Chlorophyll-*a* concentration, riverine inputs, and runoff from forestry and agricultural activities (Van Holt et al., 2012; Lara et al., 2019). In this region, upwelling activity is weaker and more intermittent than the sector equatorward from Concepción, which is dominated by strong seasonal upwelling (ca. 37S; Atkinson et al., 2002; Letelier et al., 2009; Pinochet et al., 2019).

Rocky-shore communities inhabiting the mid-intertidal zone across the province are composed of a diverse set of mobile and sessile species, where human harvesting and environmental context modulate ecological interactions (Broitman et al., 2001; Lagos et al., 2005; Caro et al., 2010; Broitman et al., 2011). The sessile communities are dominated by green macroalgae like *Ulva compressa* and *U. rigida*, red macroalgae like *Mazzaella laminarioides* and *Gelidium chilense*, or filter feeders like the barnacles *Jehlius cirratus, Notochthamalus scabrosus*, and *Perumytilus purpuratus* (reviewed in Aguilera et al., 2019).

### Experimental design and setup

At each site, we conducted a manipulative experiment with dominant-species removal as a fixed factor. The experiments were conducted between October 2014 and August 2017 in the southern region, and between December 2014 and September 2017 in the northern region—in both cases, the prolonged time span (ca. three years) allowed us to fully capture the seasonal and intra-annual variation in abiotic conditions and species composition. Prior to the start of the experiment, we estimated relative species abundances in ten 30×30 cm plots haphazardly located on the mid-intertidal zone to determine locally dominant species as those with the highest relative abundances at each site (see also Valone and Balaban-Feld, 2018). At each site, quadrats were located within an alongshore transect (ca. 3 m wide) covering ca. 50 m of the coastline. The red corticated alga *Mazzaella laminarioides* and chthamalid barnacles (a mixture of *Jehlius cirratus* and *Notochthamalus scabrosus*) were the dominant taxa in the northern region; chthamalid barnacles and the purple mussel *Perumytilus purpuratus* were the dominant species in the southern region (hereafter referred as *Mazzaella*, barnacles, and *Perumytilus*, respectively; Table 1). Before the manipulations, mean percentage cover of dominant species ranged from ca. 30 % for *Mazzaella* in the northern region to 100 % for *Perumytilus* in the southern region. Simpson’s concentration index varied between 0.4 and 0.9 in the northern and southern region, respectively, picturing a higher evenness in species abundance distribution in the former than the latter region (Table 1).

Following site characterization, we started identical field experiments. We used 30 x 30 cm experimental plots on the mid-intertidal zone of each site, which were permanently marked with stainless-steel bolts. The size of the experimental unit was selected to minimise the impact on the local intertidal communities. For logistic reasons, we deployed 10 replicate plots in the sites located on the northern sector and 20 replicates in the southern sites. Experimental plots were haphazardly located within areas of high abundance of the locally dominant species and relatively flat and horizontal rocky surfaces. Thus, we avoided crevices, tide-pools, and vertical surfaces to reduce biotic variation related to local spatial heterogeneity.

At the onset of experimental manipulations (i.e., October and December 2014 in the southern and northern regions, respectively), we haphazardly selected half of the plots at each site and completely removed the dominant species with scrapers and chisels. The remaining half of the experimental units remained un-manipulated and assigned to the control level. Settlers and recruits of the dominant species were further removed approximately every three months, i.e. the press disturbance treatment (see also Bulleri et al., 2012). Prior field studies on recruitment patterns in the province suggest that the three-month window between manipulations allows to functionally exclude dominant species (Aguilera et al., 2013; Valdivia et al., 2013; Valdivia et al., 2015).

All plots were sampled immediately before the removal of local dominants, 1-2 months after the manipulation, and then approximately every three months until the end of the experiment. All macroalgae and sessile macro-invertebrates (> 5 mm) occurring in each plot were identified *in situ* and classified to the lowest possible taxonomic level (usually species). For each plot, we used a 0.09-m^2^ frame, divided in 25 equal fields with a monofilament line, to estimate sessile species abundances as percentage cover (1 % resolution). We quantified the organisms growing on both primary and secondary substrata (i.e., rock and other organisms, respectively), which allowed total community cover to increase beyond 100 %. This protocol has been used in several studies of benthic diversity along Chile and elsewhere (see Broitman et al., 2011; Bulleri et al., 2012). The data obtained during the 11 successive surveys, conducted after the first removal, were used to estimate four dimensions of stability, i.e., resistance, resilience, recovery, and invariability. Since we were interested in the response of the remaining community to the local extinction of dominants, the latter were removed from the dataset when we assessed community stability.

### Species compensation

The degree of species compensation was measured by assessing community-wide synchrony in species abundances. To this aim, we used the *φ* statistic (Loreau and de Mazancourt, 2008):

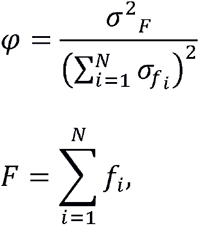

where *f_i_* is the percentage cover of species *i*, *F* is the equivalent aggregate community-level function (total community cover), and 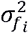 and 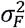 are their respective temporal variances. The index is standardised between 0 (perfect asynchrony, i.e., most species’ abundances are negatively correlated over time) and 1 (perfect synchrony, i.e., most species’ abundances are positively correlated). Since *φ* is independent of the magnitude and distribution of species abundances and variances, it allows us to quantitatively compare communities with different numbers of species (Loreau and de Mazancourt, 2008).

### Stability dimensions

Stability was assessed according to the metrics for resistance, resilience, recovery, and invariability described by Hillebrand et al. (2018). Each metric was estimated by randomly pairing one control and one disturbed plot in each site. In this way, we obtained replicate estimations within each site. The stability metrics in the functional domain were calculated on the basis of total community cover (*F*), and the stability metrics in the compositional domain were calculated on the basis of Bray-Curtis similarities (*sim*) between the disturbed and control plots (*C*).

*Resistance* (*a*) was estimated for function as the log-response ratio (*LRR*) calculated after the disturbance took place (1-2 months after the removal of local dominants):

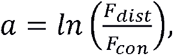

where *F_dist_* is total community cover in the dominant-removed plot, and *F_con_* is the same property in the control plot. A value of *a* = 0 indicates maximum resistance of the community to the disturbance, *a* < 0 indicates low resistance due to underperformance, and *a* > 0 low resistance due to overperformance relative to controls, respectively (Hillebrand et al. 2018). This means that both initially increased and decreased functioning will reflect low functional resistance. For that reason, we used the absolute value of resistance in some hypothesis tests as described below (see *Statistical analyses: H1 and H2*).

The resistance of composition (*a*) was calculated on the basis of Bray-Curtis similarities between the disturbed and the control plots:

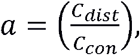

Where *C_dist_* and *C_con_* are the species-abundance matrices in the dominant-removed and control plots, respectively. Maximum similarities of *a* = 1 indicate 100 % of similarity between disturbed and control plots and thus maximum resistance. Minimum similarities of *a* = 0 indicate 0% of similarity between both conditions and thus extremely low resistance.

*Resilience* (*b*) was calculated for each plot as the slope of the linear regression of *LRR* calculated for each sampling event (Hillebrand et al. 2018):

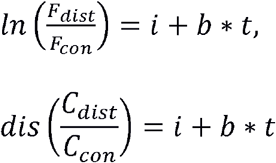

where *i, b*, and *t* are the intercept, slope, and time, respectively. A *b*-value equalling zero indicates no recuperation over time, *b* > 0 indicates faster recuperation in case of a negative initial effect of disturbance on the function or composition, and *b* < 0 indicates further deviation from control in case of negative initial effects. When the disturbance generates positive effects on functioning (i.e., *a* > 0), *b* > 0 is interpreted as further deviation from control and vice versa.

*Recovery* (*c*) was estimated as the *LRR* and similarity calculated at the end of the experiment, *c*-values equal to zero denote maximum recovery, c < 0 incomplete recovery, and c > 0 overcompensation relative to control. Like for resistance, both c < 0 and c > 0 reflect low functional recovery; thus, the absolute value of functional *c* was analysed in the test of hypotheses H1 and H3 (see *Statistical analyses*).

*Invariability* (*d*) was calculated as the multiplicative inverse of the standard deviation of residuals (*resid*) from the regression slope *b* as:

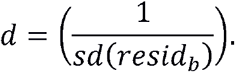

Larger values de *d* denote smaller temporal variations around the trend of recovery in function or composition (Hillebrand et al. 2018). Note that invariability is termed as *Temporal stability* by Hillebrand et al. (2018). However, we have preferred to use *Invariability* to avoid confusions with the term *Stability* (see also Radchuk et al., 2019).

### Statistical analyses

For simplicity, hereafter the community in each site will be referred to according to the identity of the locally dominant species that was removed (i.e., *Mazzaella*-, barnacle-, or *Perumytilus*-dominated communities). This means that the plots and sites will be grouped into each of the three community types in some of the statistical analyses.

### H1: The magnitude of the stability response will depend on the identity of the removed dominant species

To test this hypothesis, we first used *t*-tests for each community and within each stability domain (either functional or compositional) to determine if resistance, resilience, and recovery significantly differed from the reference values. Secondly, we used separate general lineal models (LMM) to compare the magnitude of each stability dimension among the three communities. Accordingly, we used the absolute value of functional resistance and recovery to focus on their magnitudes. The models included the identity of the removed dominant species as fixed effect, and site as random effect. Nevertheless, site was removed from the models of the compositional domain due to singularity (i.e. variance nearly zero).

### H2: Across sites, the removal of the dominant species will elicit negative or neutral correlations between functional and compositional stability dimensions due to prevailing compensatory dynamics

First, we used one-tailed t-tests to determine whether *φ* was significantly lower than 1 (i.e., perfect synchrony in species fluctuations) for each community. In addition, we used a generalised linear mixed model (GLMM) to determine if the removal of the locally dominant species decreased *φ* (i.e., increased the degree of species compensation within the community). For the model we used a Gamma structure of errors and a log link function; the model included the identity of the dominant species in each site and the removal treatment (removal or control) as crossed and fixed factors, and the site (eight levels) as random factor. After initial the fit, however, site was removed due to singularity.

Second, Pearson product-moment correlations were calculated between functional and compositional dimensions of stability. Correlations were calculated separately for each community and stability dimension. In these correlations, we used the absolute value of every functional stability dimension to assess the magnitude, rather than the direction, of these responses. The absence of statistically significant correlations, in addition to near-zero values of *φ* were used as evidence of species compensation after the removal of dominant species.

Third, to identify the species driving compensation (or the lack thereof), we used Similarity Percentage Routines (SIMPER; Clarke, 1993) that partitioned the contribution of each species to the between-treatment dissimilarity in each community.

### H3: Across sites and within the stability domains (functional and compositional), the stability dimensions will be correlated with each other

Pearson product-moment correlations were used to test this hypothesis. Within the functional and compositional domains, analyses were conducted combining all sites and communities to test for a tendency between these dimensions to correlate independently of the ecological context and species identities.

All analyses, calculations, and plots were done in R programming environment version 4.0.2 and the R-packages *lme4, MuMIn, sjPlot, tidyverse*, and *vegan* (Bates et al., 2015; Oksanen et al., 2019; Wickham et al., 2019; Bartoń, 2020; Lüdecke, 2020; R Core Team, 2020), except for Fig. 1, which was made using MatLab R2014a and data from MODIS-Aqua.

## Results

### H1: The magnitude of the stability response will depend on the identity of the removed dominant species

The different communities significantly responded to the experimental disturbances within both functional and compositional domains (Figs. 2A, C, E, G and B, D, F, H, respectively; Table A1 in the Appendix). The exception was the functional resilience of the barnacle- and *Perumytilus*-dominated communities, which did not differ from zero (Table A1, Fig. 2C). When we compared the magnitude of the responses among communities, the three communities exhibited remarkably similar stability responses to the experimental disturbance for both domains (Fig. 2). Accordingly, we did not detect statistically significant differences among the communities, with the exception of functional resilience (GLMM contrasts in Table A2: *Mazzaella* > barnacles; Fig. 2C).

**Fig. 2.**
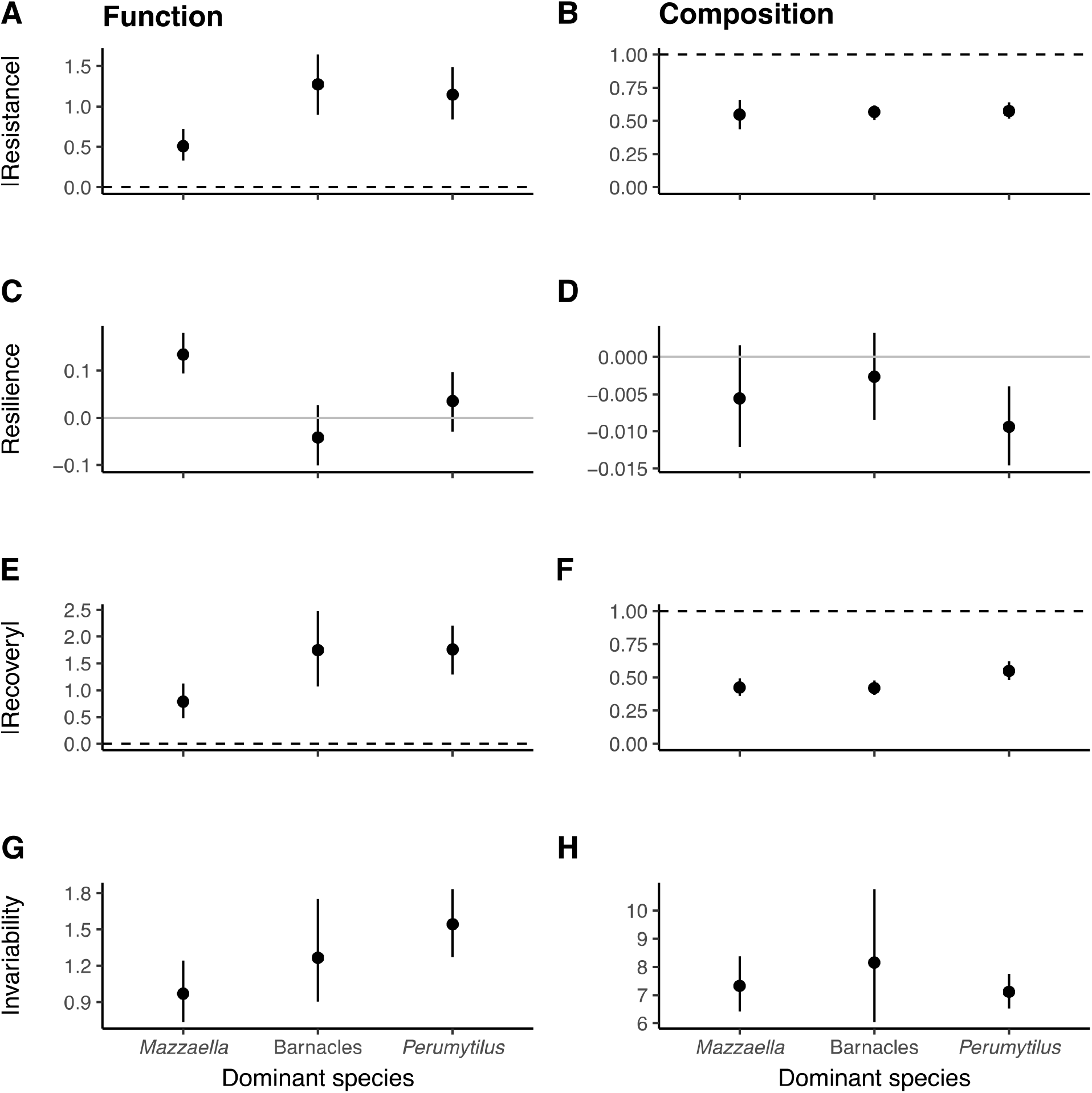
Four dimensions of stability in the (A, C, E, G) functional and (B, D, F, H) compositional domains. Note that (A) functional resistance and (E) recovery are shown as absolute values in order to compare magnitudes, rather than directions of effects. Values are mean ± 95 % confidence intervals. Dashed lines represent maximum resistance and recovery; solid grey lines represent minimum resilience. All stability dimensions statistically differed from reference values, excepting (C) functional resilience of barnacle- and *Perumytilus*-dominated communities (t-tests: *t_16_* = −1.24, *P* > 0.05; *t_28_* = 1.28, *P* > 0.05, respectively). Accordingly, the three types of communities exhibited similar stability responses, excepting functional resilience (GLMM contrasts: *Mazzaella* > barnacles; *Mazzaella* > *Perumytilus*).

### H2: Across sites, the removal of the dominant species will elicit negative or neutral correlations between functional and compositional stability dimensions due to prevailing compensatory dynamics

The low values of the proxy for species synchrony, *φ*, indicated that species compensation was common across communities and removal treatments (Fig. 3). The parameter was on average 0.11 (0.06 – 0.16), 0.30 (0.21 – 0.40), and 0.17 (0.12 – 0.21) for *Mazzaella*, barnacles, and *Perumytilus* communities respectively (mean [95% CI]). These values were significantly lower than 1 in each community (one-tailed *t*-tests: *t_9_* = −42.2, *t_16_* = −17.5, and *t_28_* = −38.2 for *Mazzaella*, barnacles, and *Perumytilus*, respectively; *P* < 0.001). However, the response of these dynamics to the dominant removal varied among communities. The press disturbance increased the strength of species compensation in the *Perumytilus* communities, but not in the others (GLM: *R^2^* = 0.45, *P* < 0.001 for an 80-% decrease in *φ* in the *Perumytilus* communities, see Figure 3). In these communities, the strengthening of species compensation was accounted for by the overshoot in the abundance of Ulvoids (mixture of *Ulva rigida* and *U. compressa*) during the first half of the experiment (late 2014 − early 2016; see Fig. A1 in the Appendix); this taxon was replaced by chthamalid barnacles over time, which remained relatively stable until the end of the experiment in mid-2017 (Fig. A1).

**Fig. 3.**
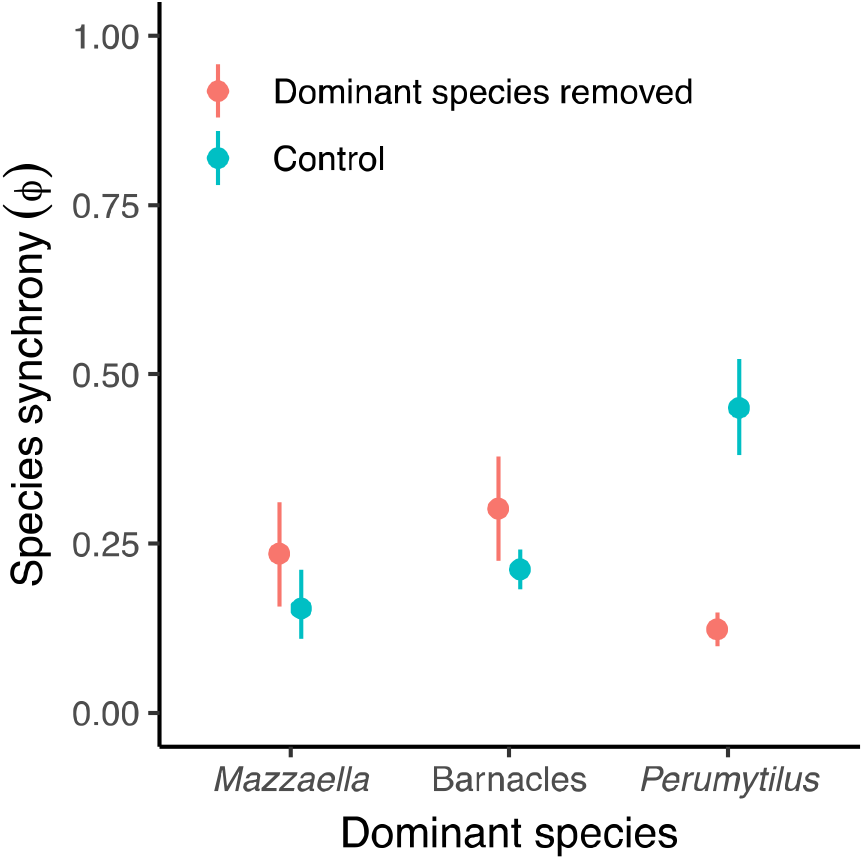
Community-wide synchrony among subordinate species in each community type (*Mazzaella*-, barnacle-, or *Perumytilus*-dominated community). The range of *φ* is [0,1], with 1 = perfectly synchronous species fluctuations and 0 = perfectly asynchronous, compensatory species fluctuations. Values are mean ± 95 % confidence intervals. All communities statistically differed from 1 (one-tailed *t*-tests: *t_9_* = −42.2, *P* < 0.001; *t_16_* = −17.5, *P* < 0.001; *t_28_* = −38.2, *P* < 0.001, respectively), indicating prevalent compensatory dynamics across the region. The removal of the dominant species systematically increased the strength of species compensation in the *Perumytilus* communities, but not in the others (Gamma GLM, *R^2^* = 0.45, P < 0.001 for *Perumytilus*).

About seven (TEMB and GUAN) and ten (PTAL) taxa accounted for ca. 90 % of the dissimilarities between treatments in the northern region; about three (CALF) and five (CHAI) taxa explained the 90 % of the multivariate differences between treatments in the southern metacommunity (Table A3). Across both regions, these species included encrusting, filamentous, and corticated macroalgae, in addition to barnacles (e.g., *Austromegabalanus psittacus*; Table A3). The lichen *Verrucaria* sp. accounted for a relatively important percentage of between-treatment dissimilarities in the northern region (TEMB, GUAN, and PTAL; Table A3).

According to the high prevalence of species compensation, we observed almost no statistically significant correlation between functional and compositional stabilities across communities (Fig. 4). Separately for the three community types, we observed non-statistically significant, correlations for resistance (Fig. 4A; Table A4 in the Appendix). For resilience, only the communities in which barnacles were removed showed a significant, and negative, correlation between the functional (absolute value) and compositional domains (green symbols in Fig. 4B; Table A4). Similarly, the barnacle-dominated communities exhibited a negative correlation between functional (absolute value) and compositional recoveries (green symbols in Fig. 4C; Table A4). Finally, functional invariability was uncorrelated from compositional invariability (Fig. 4D; Table A4).

**Fig. 4.**
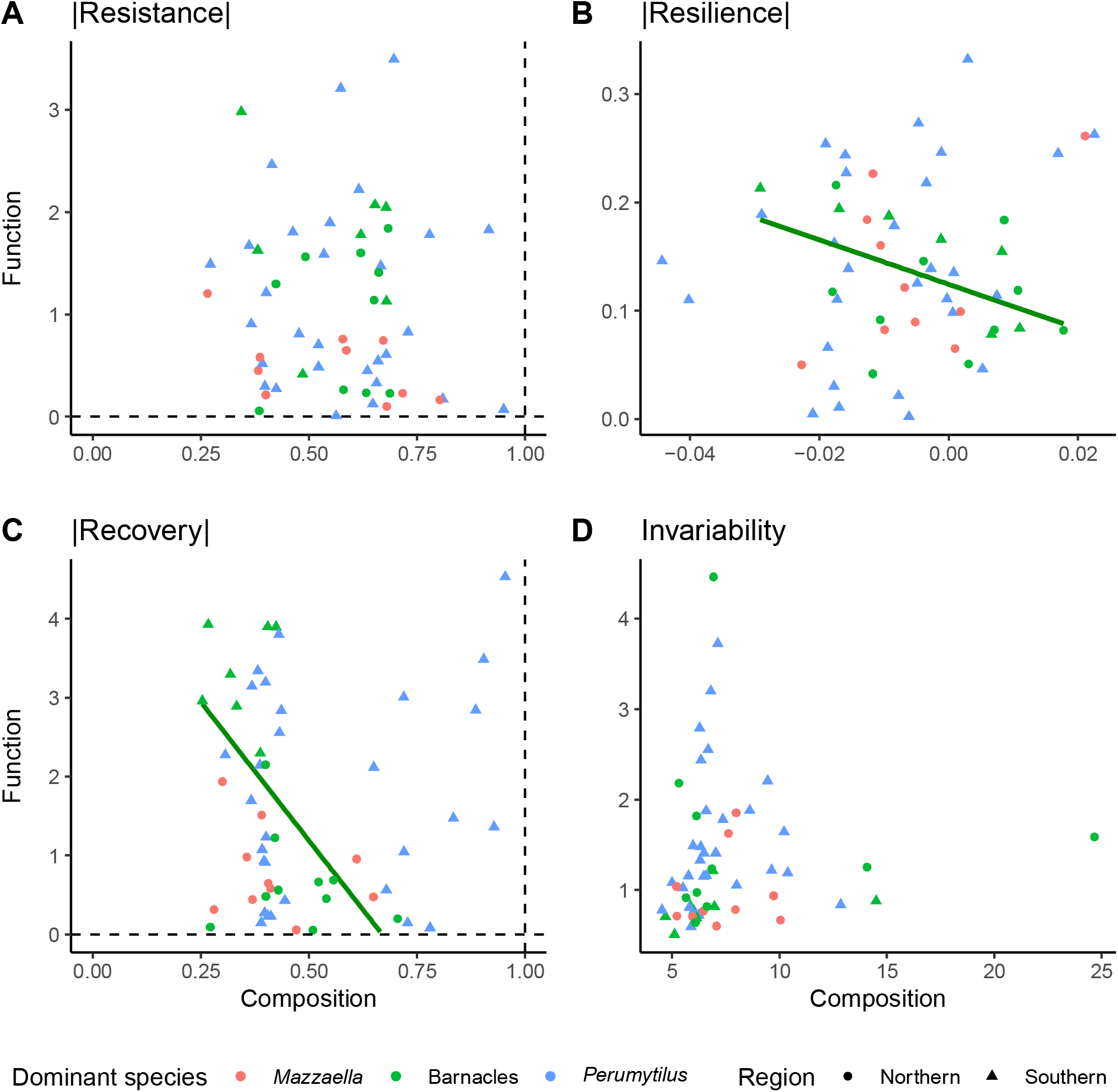
Pairwise associations between functional and compositional dimensions of stability. In these analyses, all functional dimensions were used as absolute values in order to correlated magnitudes, rather that directions of effects. Dashed lines represent maximum resistance and recovery; solid grey lines represent minimum resilience. Only (B) resilience and (C) recovery of the barnacle-dominated communities (green symbols and trend lines ± 95 % confidence intervals) showed significant function-composition correlations (r = −0.52 and r = −0.57, respectively; P < 0.05).

### H3: Across sites and within the stability domains (functional and compositional), the stability dimensions will be correlated with each other

Our results did not support this hypothesis, yet we detected some significant correlations (Fig. 5; Table A5 in the Appendix). In the functional domain, statistically significant correlations were found between resistance and resilience (negative; Fig. 5A; Table A5), resistance and invariability (positive; Fig. 5D; Table A5), and resilience and invariability (negative; Fig. 5E; Table A3). In the compositional domain, we did not detect statistically significant correlations among stability dimensions (Fig. 6; Table A5).

**Fig. 5.**
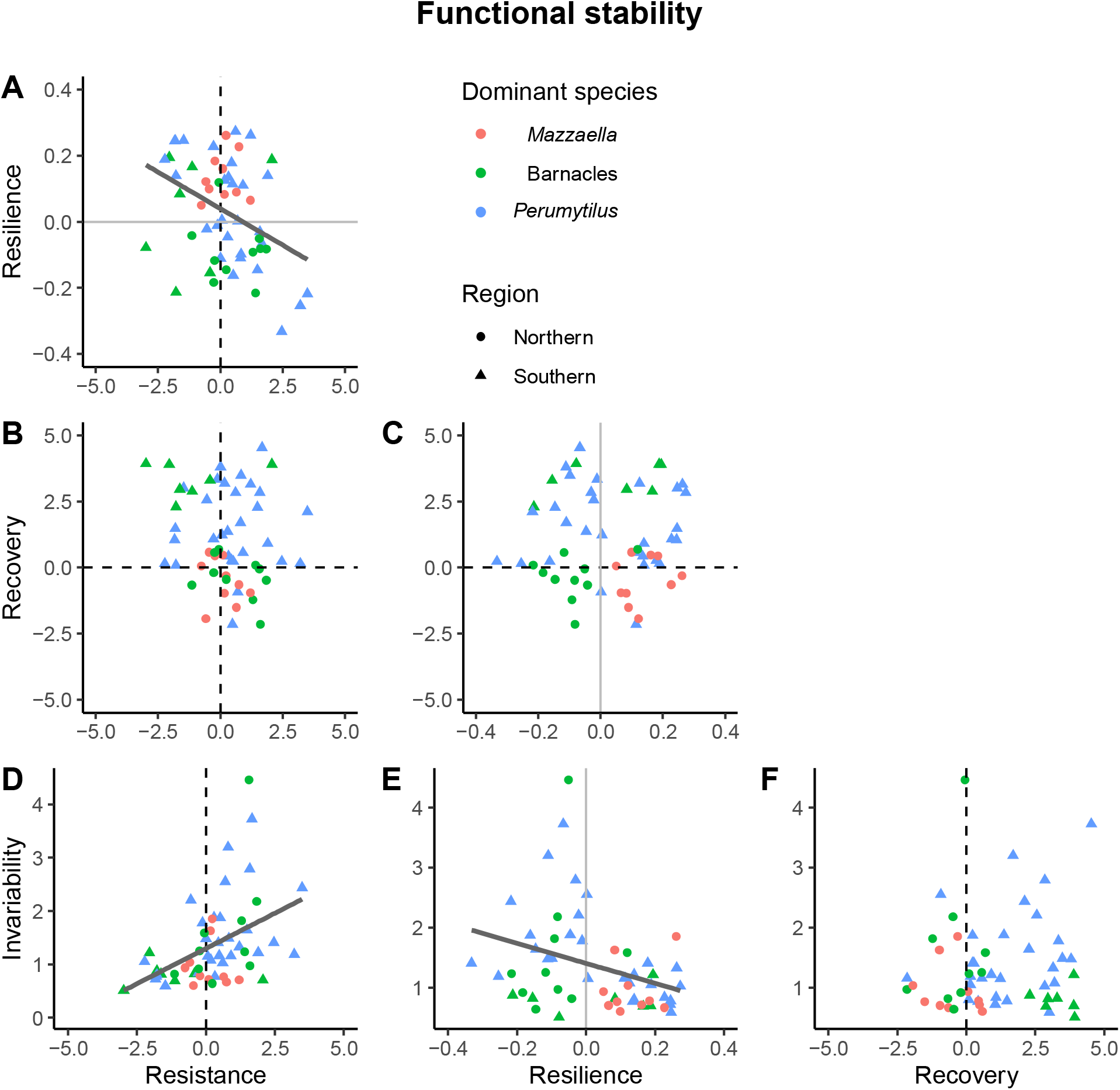
Associations among four dimensions of functional stability. Dashed lines represent reference values for maximum resistance and recovery; solid grey lines represent reference values for minimum resilience. Across communities, statistically significant correlations were detected for the resistance-resilience, resistance-invariability, and resilience-invariability pairs (trend lines ± 95 % confidence intervals; r = −0.38, r = 0.39, and r = −0.36, respectively; P < 0.01).

**Fig. 6.**
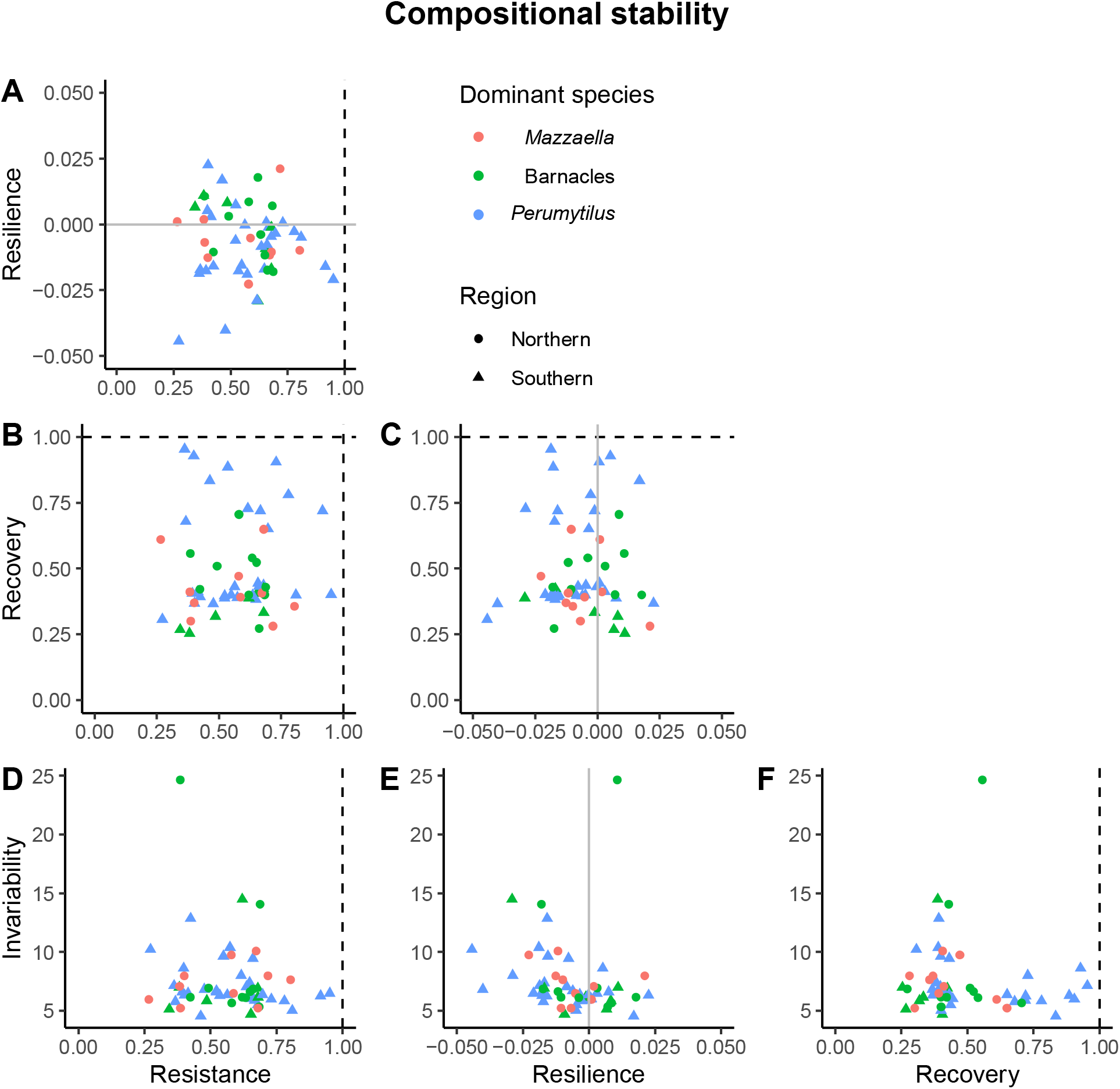
Associations among four dimensions of compositional stability. Dashed lines represent maximum resistance and recovery; solid grey lines represent minimum resilience. No statistically significant correlation was detected across communities.

## Discussion

This study showed that the stability of rocky intertidal communities in a broad biogeographical province of the SE Pacific shore is a multidimensional property. After the removal of locally dominant species, the magnitude of the effect of the disturbance and the temporal pattern of recovery were surprisingly similar among communities but varied between functional and compositional response variables, and among the four dimensions of stability examined here; namely, resistance, resilience, recovery, and invariability. Only in the functional domain we still observed significant correlations between resistance and resilience (negative), resistance and invariability (positive), and resilience and invariability (negative). The high prevalence of species compensation and asynchronous dynamics could explain the general lack of correlation between functional and compositional stability domains, indicating that functional recovery may not involve a fully compositional recovery in this study system. Our results are in line with recent evidence of the multifaceted nature of ecological stability (Donohue et al., 2016; Hillebrand et al., 2018; Pennekamp et al., 2018; Kéfi et al., 2019; Radchuk et al., 2019), which we further discuss below.

### Different communities exhibited similar initial and long-term stability responses to local extinction

The disturbance elicited significant stability responses within both the functional and compositional domain regardless of the dominant species identity, with the exception of functional resilience, which was indistinguishable from zero in the case of the invertebrate-dominated communities. Despite these differences, no community fully recovered in the end of the experiment, 35 months after the start of the disturbance. Importantly, the consistently negative compositional resilience provides further support for a persistent departure of these assemblages from the reference state (i.e. control plots), as shown in phytoplankton mesocosm experiments (Hillebrand et al., 2018). This result may indicate that the recovery, and restauration, of this ecosystem may need a longer time span than here considered. Long-term (i.e., 20 years) monitoring of rocky intertidal communities suggests the existence of cyclic succession after severe disturbances, with the alternation of dominant species every ca. two years (Benincà et al., 2015). These nonequilibrium dynamics can well be amplified by environmental stochasticity, as shown by experimental removals of dominant mussels (Wootton, 2010). Thus, the non-equilibrium and near-chaotic nature of coastal systems may obstruct our ability to restore disturbed coastal ecosystems. Indeed, a major review of restoration strategies indicates that functional and compositional recovery of coastal systems may reach on average a 34 % in 12 years, even when active restoration is enforced (Moreno-Mateos et al., 2020).

Between-site environmental differences could have generated “unexplained” variability in the stability responses, precluding our ability to detect differences among the dominant species. Nevertheless, our analyses suggest that the factors (other than dominant’s identity) associated to between-site differences may have little influence on the stability measurements (compare the pseudo-*R^2^* between fixed factors and full models in Table A2). In the case of functional stability, between-site differences accounted for small, but non-zero, amount of variation in the stability responses. Dampened environmental variability in sites under the influence of persistent upwelling could have influenced the invariability in community cover (Valdivia et al., 2013). Sea water temperature patterns define the phenology of most intertidal organisms (Sanford, 2002; Baldanzi et al., 2018), which, mediated through larval dispersal processes, can influence overall population growth rates, and thus functional resilience and recovery (Svensson et al., 2005). In the case of compositional stability, on the contrary, site accounted for almost no variation in Bray-Curtis dissimilarity. If site differences seemly accounted for a small variation in stability responses, then, to what degree the stability observed at the local scales can scale up to regional and seascape scales?

The area-invariability relationship indicates that stability should increase from local to regional scales due to asynchronous dynamics among sites within the landscape, and also among species within sites (Wang and Loreau, 2014; Zhang et al., 2018). Moreover, increased beta diversity (i.e. between-site differences in species composition) is expected to increase the spatial asynchrony in functioning among sites, and thus the stability of the whole landscape (Wang and Loreau, 2016). According to the high prevalence of asynchronous species dynamics (this study; Valdivia et al., 2013) and high beta diversity (Broitman et al., 2001; Valdivia et al., 2015; Valdivia et al., 2017), predictions based on the area-invariability relationship should hold in our study system. Albeit the extension of this theory to other stability dimensions and the compositional domain still needs further development, we suggest that maximising protected areas, rather than focusing on patches that contain unique assemblages, should be a conservation goal to maintain the functioning in the analysed ecosystem (see also Economo, 2011; Ospina-Alvarez et al., 2020).

### Compensatory dynamics were common within the system, leading to null or negative function-composition correlations

In agreement with our prediction, asynchronous and compensatory species dynamics were strong in the study region. The strength of these dynamics, however, varied among the three analysed community types. In particular, the removal of *Perumytilus* reduced the synchrony in species abundances following compensatory dynamics between ulvoid algae and barnacles, according to the facilitation model of ecological succession (Connell and Slatyer, 1977). In addition to competitive species replacement during succession, asynchronous species dynamics can also reflect differential species environmental responses, population compensatory cycles, species-wise variations in dispersal, and neutral, zero-sum dynamics (e.g. Lehman and Tilman, 2000; Chesson et al., 2001; Loreau et al., 2003; Loreau and de Mazancourt, 2008). At global scales, asynchronous dynamics can have stronger effects on stability than species richness (Valencia et al., 2020). Accordingly, evidence of the central role of species synchrony in the stability of ecosystem functions is supported by studies across ecosystems and taxonomic groups (reviewer by González and Loreau, 2009).

The lack of correlation between functional and compositional measurements of stability could well reflect the effects of species compensation. For instance, high resistance and recovery (i.e., near-zero log-response ratios) in the functional domain were observed along ample ranges of both metrics in the compositional domain, and the negative function-composition correlations detected for resilience and recovery further reinforce the notion of functional compensation within these communities (Guelzow et al., 2017; Hillebrand et al., 2018; Hillebrand and Kunze, 2020). This indicates that a given level of functioning could be performed by communities with different compositions, which may have important consequences for ecosystem restauration (Fig. 7A; Borja et al., 2010).

**Fig. 7.**
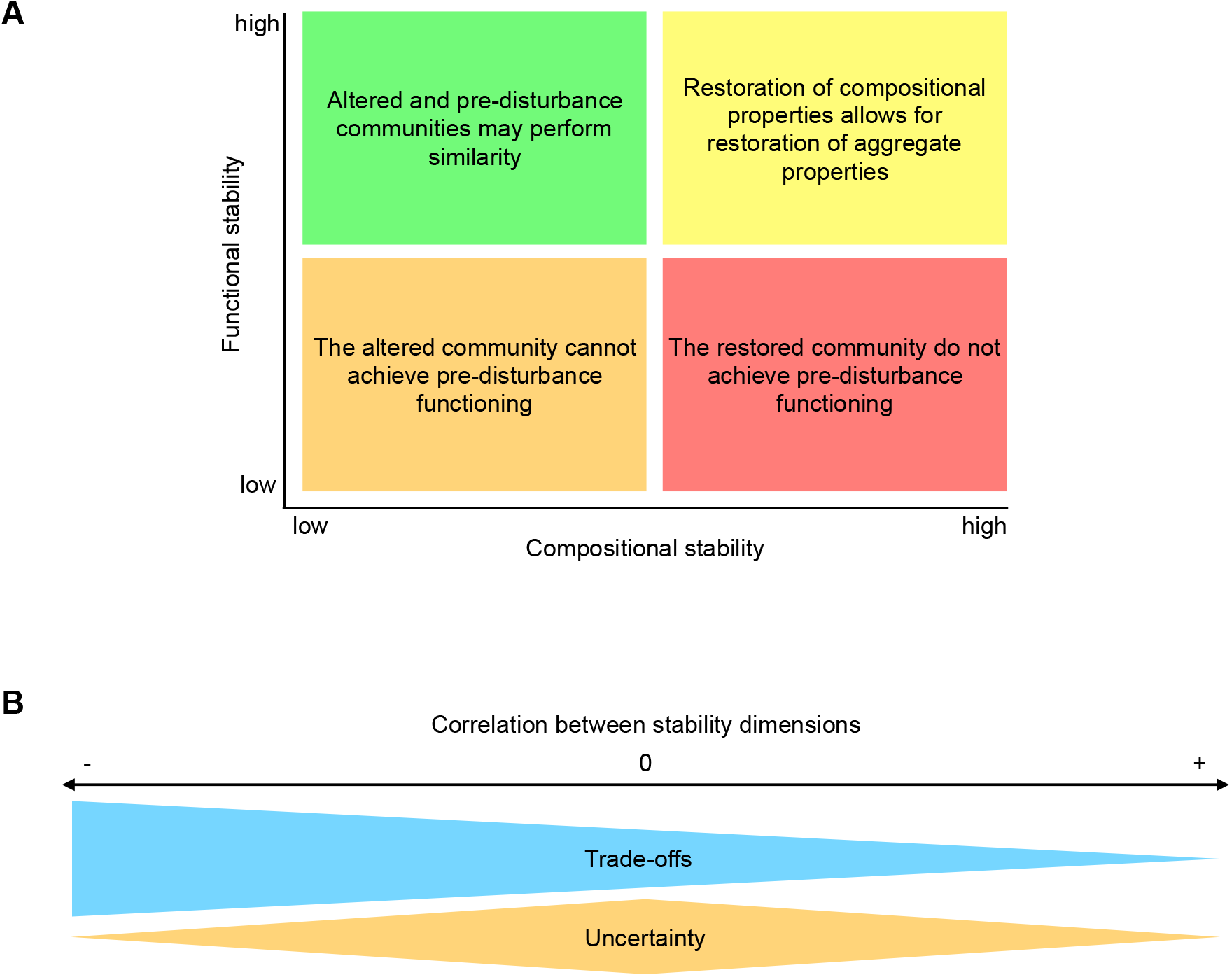
Summary of scenarios based on correlations between (A) functional and compositional stability domains and (B) stability dimensions. Different scenarios have different scenarios for ecosystem conservation and management. Panel A: Systems exhibiting positive functional-compositional stability correlations may need a full recovery in species composition to achieve a full recovery in functioning (yellow quadrant); if not, no functional recovery would be possible (orange quadrant). Contrarily, negative functional-compositional correlations (e.g., barnacle communities in this study) imply that the altered community may be able to compensate the functioning of the pre-disturbance community, but also that restored communities are unable to recover pre-disturbance functional values. Panel B: A positive correlation between two aspects of stability (e.g., functional resistance and invariability; this study) indicates that and conservation decisions could improve both aspects of stability; a negative correlation (e.g., functional resistance and resilience; this study) implies that policy trade-offs will be crucial to maintain either one or another aspect of stability. The lack of correlation among stability dimensions (e.g., all compositional dimensions in this study) indicates that, due to high uncertainty, multiple aspects are mandatory to characterise the stability of the ecosystem.

### The stability dimensions were mostly uncorrelated across community types and stability domains

In the functional domain, the correlations among the different dimensions of stability ranged from negative to null and to positive. We were able to detect negative correlations between functional resistance and resilience and between resilience and invariability. Negative functional resistance-resilience and resilience-invariability correlations have been also observed in modelling and experiments conducted on experimental phytoplanktonic mesocosm communities (Hillebrand et al., 2018), and have been associated to comparatively high growth rates that allow faster recovery rates with stronger initial impacts (Harrison, 1979). Similarly to aquatic ciliate communities (Pennekamp et al., 2018), larger initial functional responses (i.e. lower resistance) were associated with more temporally stable functioning, which could respond to a positive relationship between species’ invariability and relative abundance (Grman et al., 2010; Lamy et al., 2020).

In the compositional domain, on the other hand, no statistically significant correlation between stability dimensions was detected. This high dimensionality implies that the four aspects of compositional stability analysed in this study are fully orthogonal and are mandatory to completely describe the response of these communities to an ecologically realistic disturbance akin to our press removal of the dominant species. Uncorrelated stability dimensions have been highlighted by previous work when ecological communities face severe environmental disturbances (Allison, 2004; van Ruijven and Berendse, 2010; Donohue et al., 2013; Hillebrand et al., 2018). Contrarily, most empirical research has been based on one or two stability dimensions; the temporal coefficient of variation has been widely used to assess the stability in aggregate ecosystem functions in the absence of major disturbances (reviewed by Kéfi et al., 2019). Yet, restoration assessment after pulse and press disturbances would largely benefit from considering multiple components of stability, because, as we have shown, these components may well be decoupled during the process of recovery (Fig. 7B; Donohue et al., 2016; Moreno-Mateos et al., 2020).

Perhaps, percentage cover would have been too coarse to detect differences stability among community types and positive correlations among the stability dimensions. Albeit we considered the multi-layered nature of macrobenthic communities in our in-situ estimations, it would be the case that between-plot variations in the tri-dimensional projection of the communities would have generated unwanted variation in the stability responses. However, our results are well supported by previous work assessing other functional proxy, such as consumption rates and biomass (Ghedini et al., 2015; Hillebrand et al., 2018; Valencia et al., 2020). Percentage cover in sessile communities is the results of the complex interaction between preemptive competition, disturbance, and recolonisation (Paine, 1980), which makes it a good proxy for functioning.

### Conclusion and future directions

Our results allow us to conclude that ecological stability of the analysed intertidal communities is a multidimensional and complex phenomenon. The experimental removal of locally dominant species elicited significant, and similar in magnitude, stability responses of communities in a large biogeographical province of the SE Pacific shore. These responses varied between functional and compositional domains, likely due to pervasive and strong asynchronous species dynamics.

Still is necessary to determine how different types and aspects of disturbances can influence stability dimensionality in these ecosystems. Albeit recent modelling work indicates that disturbance types such as random, species-specific, or local disturbances can have similar effects on stability dimensionality (Radchuk et al., 2019), contrasting the effects of pulse and press disturbances can enhance our mechanistic understanding of the ecological consequences of current climate crisis (Kéfi et al., 2019)—anthropogenic climate change involves the interdependent effects of both types of disturbances over multiple spatiotemporal scales (Harris et al., 2018). Regional manipulative experiments considering a more ample range of environmental filters and dispersal limitations would allow us model the role of spatial aspects of disturbance (Zelnik et al., 2018; 2019) in how stability dimensionality scales from sites to entire regions; to our best knowledge, the scale dependence of stability has been investigated only for single aspects of stability (Wang et al., 2017). In summary, realising that stability is indeed a multidimensional concept will enhance our ability to predict the response of communities and metacommunities to future and ongoing anthropogenic disturbances.

## Acknowledgements

We thank all the enthusiastic undergraduate interns (Biología Marina; UACH), members of the Coastal Ecology Lab (Lafkenchelab, UACH), and the Changolab (CEAZA-UAI) who provided invaluable help during the fieldwork. Melissa González, José Pantoja, and Oscar Beytía were fundamental to complete the sampling schedule in the northern region. Constructive criticism by Fabio Bulleri, Matt Bracken and three anonymous reviewers greatly improved an early version of this manuscript. This study was financially supported by the FONDECYT #1190529, FONDECYT #1141037, and FONDAP #15150003 (IDEAL) to NV. MAA was supported by FONDECYT #1160223 and PAI-CONICYT #79150002-2016. BRB acknowledges the support of FONDECYT #1181300 and the MUSEL Millennium Nucleus, funded by Iniciativa Científica Milenio.

## Author contribution

NV and BRB conceived the study; NV analysed the data; BRB and MAA contributed with field work, reagents and analytical material; NV wrote the paper; BRB and MAA contributed significantly to the writing of the manuscript.

## Conflicts of interest

The authors have no conflict of interest.

Permit(s) – The study did not need permits, because all experiments were conducted on open-access coastal areas.

## Notes

### Competing Interest Statement

The authors have declared no competing interest.

