## Supplemental Figure and Tables for "High dimensionality of the stability of a marine benthic ecosystem"

^2^Centro FONDAP de Investigación de Dinámicas de Ecosistemas Marinos de Altas Latitudes (IDEAL)

^3^Departamento de Biología Marina, Facultad de Ciencias del Mar, Universidad Católica del Norte, Larrondo 1281, Coquimbo, Chile

^4^Centro de Estudios Avanzados en Zonas Áridas (CEAZA), Universidad Católica del Norte, Ossandón 877, Coquimbo, Chile

^5^Departamento de Ciencias, Facultad de Artes Liberales & Bioengineering Innovation Center, Facultad de Ingeniería y Ciencias, Universidad Adolfo Ibáñez, Av. Padre Hurtado 750, Viña del Mar, Chile

Nelson Valdivia ORCID ID: https://orcid.org/0000-0002-5394-2072

Bernardo R. Broitman ORCID ID: http://orcid.org/0000-0001-6582-3188

Moisés A. Aguilera ORCID ID: https://orcid.org/0000-0002-3517-6255

Table A1. Statistical tests (*t*-statistic from t-tests) of resistance, resilience, and recovery against reference values. The modulus of functional resistance and recovery are used to assess the magnitude of these stability responses. Functional resistance and recovery were tested against 0 (maximum value). Compositional resistance and recovery were tested against 1 (maximum value). Both, functional and compositional resilience were tested against 0 (minimum resilience).

| Community | DF | Function | | | | | | Composition | | | | | |
| --- | --- | --- | --- | --- | --- | --- | --- | --- | --- | --- | --- | --- | --- |
|  |  | \|Resistance\| | | Resilience | | \|Recovery\| | | Resistance | | Resilience | | Recovery | |
| *Mazzaella* | 9 | 4.83 | ******* | 6.18 | ******* | 4.31 | ******* | -8.21 | ******* | -262.34 | ******* | -15.00 | ******* |
| Barnacles | 16 | 6.23 | ******* | -1.24 |  | 4.87 | ******* | -15.01 | ******* | -313.01 | ******* | -19.97 | ******* |
| *Perumytilus* | 28 | 6.47 | ******* | 1.28 |  | 7.05 | ******* | -13.46 | ******* | -366.08 | ******* | -11.53 | ******* |
| ‘***’ *P* < 0.001 | | | | | | | |  |  |  |  |  |  |

Table A2. Summary of LMMs for the effect of the identity of the removed dominant species (*Mazzaella*, barnacles, or *Perumytilus*) on the magnitude of stability responses. The modulus of functional resistance and recovery are used in these analyses. *R^2^_m_* and *R^2^_c_* are marginal and conditional pseudo coefficients of determination that express the proportion accounted for the fixed (dominant species identity) factors and entire model (species identity and site), respectively. For the compositional response variables, the random factor accounted nearly zero variance.

| Domain | Dimension | *R^2^_m_* | *R^2^_c_* | Significant contrast |  |
| --- | --- | --- | --- | --- | --- |
| Functional | \|Resistance\| | 0.09 | 0.19 |  |  |
|  | Resilience | 0.19 | 0.46 | *Mazzaella* > barnacles | * |
|  | \|Recovery\| | 0.08 | 0.38 |  |  |
|  | Invariability | 0.09 | 0.19 |  |  |
| Compositional | Resistance | 0.01 | 0.01 |  |  |
|  | Resilience | 0.04 | 0.04 |  |  |
|  | Recovery | 0.12 | 0.12 |  |  |
|  | Invariability | 0.02 | 0.02 |  |  |
| ‘*’ *P* < 0.05 | | | | |  |

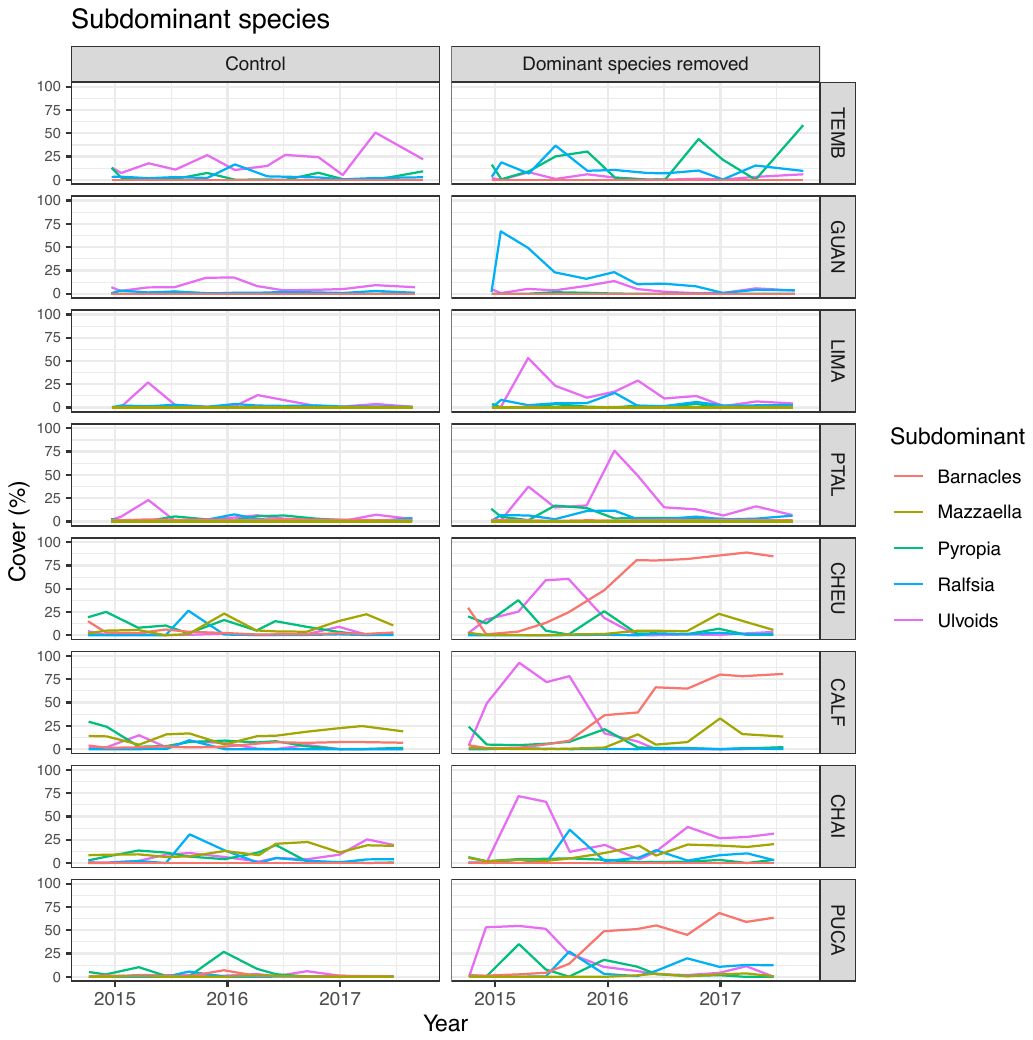

Fig. A1. Temporal patterns of subdominant taxa in control (dominant-present) and disturbed (dominant-removed) experimental plots (left and right panels, respectively) at each rocky intertidal site. Barnacles = mixture of the chthamalid barnacles *Jehlius cirratus* and *Notochthamalus scabrosus;* *Mazzaella* = *M. laminarioides;* *Pyropia* = *Pyropia* spp.; *Ralfsia* = *R. verrucosa*; Ulvoids = mixture of *Ulva rigida* and *U. compressa*. TEMB, GUAN, LIMA, and PTAL were located in the northern region; CHEU, CALF, CHAI, and PUCA were located in the southern region. Only taxa accounting for > 10 % of the between-treatment dissimilarities in each site are shown (see breakdown of dissimilarities in Table A3). Removed dominant species in each site were *Mazzaella* (LIMA and PTAL), barnacles (TEMB, GUAN, and CHAI), and *Perumytilus purpuratus* (CHEU, CALF, and PUCA).

Table A3. Breakdown of mean contribution of each taxon to the differences in community structure (Bray-Curtis dissimilarities) between control and disturbed macrobenthic sessile communities across eight sites spanning a broad province. Taxa accounting for ca. 90 % of the between-treatment dissimilarities are shown. Also, we provide the mean contribution, standard deviation (sd), mean percentage cover in control and disturbed plots, and cumulative contribution of each taxon. Removed dominant species in each site is bracketed.

| **TEMB (barnacles)** |  |  |  |  |  |
| --- | --- | --- | --- | --- | --- |
| Taxon | Mean contribution | sd | Control | Disturbed | Cumulative contribution |
| Ulvoids | 0.11 | 0.12 | 4.51 | 22.11 | 0.21 |
| *Pyropia* spp. | 0.10 | 0.13 | 18.53 | 3.73 | 0.41 |
| *Polysiphonia* spp. | 0.09 | 0.10 | 1.76 | 16.13 | 0.58 |
| *Ralfsia verrucosa* | 0.07 | 0.09 | 13.05 | 4.82 | 0.71 |
| *Notobalanus flosculus* | 0.06 | 0.09 | 1.16 | 12.55 | 0.83 |
| *Austromegabalanus psittacus* | 0.03 | 0.06 | 1.00 | 7.04 | 0.89 |
| *Hildenbrandia lecannellieri* | 0.01 | 0.05 | 3.38 | 2.25 | 0.92 |
| **GUAN (barnacles)** |  |  |  |  |  |
| Taxon | Mean contribution | sd | Control | Disturbed | Cumulative contribution |
| *R. verrucosa* | 0.11 | 0.11 | 20.65 | 2.58 | 0.22 |
| *Hypnea* sp. | 0.09 | 0.10 | 1.33 | 15.75 | 0.39 |
| *Polysiphonia* spp. | 0.07 | 0.11 | 5.89 | 10.73 | 0.54 |
| *H. lecannellieri* | 0.06 | 0.09 | 10.91 | 5.15 | 0.66 |
| *Verrucaria* sp. | 0.06 | 0.05 | 11.05 | 2.20 | 0.77 |
| Ulvoids | 0.06 | 0.07 | 5.84 | 9.38 | 0.88 |
| *Gelidium* spp. | 0.02 | 0.03 | 1.22 | 3.56 | 0.92 |
| **LIMA (*Mazzaella*)** |  |  |  |  |  |
| Taxon | Mean contribution | sd | Control | Disturbed | Cumulative contribution |
| Ulvoids | 0.12 | 0.15 | 21.22 | 9.33 | 0.29 |
| *Gelidium* sp. | 0.08 | 0.07 | 11.22 | 13.25 | 0.49 |
| *Polysiphonia* spp. | 0.04 | 0.06 | 6.67 | 4.62 | 0.60 |
| *Lithothamnion* sp. | 0.04 | 0.04 | 3.80 | 8.31 | 0.70 |
| *H. lecannellieri* | 0.03 | 0.04 | 4.02 | 6.47 | 0.78 |
| *R. verrucosa* | 0.03 | 0.04 | 5.80 | 2.87 | 0.85 |
| *Corallina officinalis* | 0.02 | 0.04 | 2.27 | 3.33 | 0.89 |
| *Verrucaria* sp. | 0.01 | 0.01 | 2.85 | 2.25 | 0.92 |
| **PTAL (*Mazzaella*)** |  |  |  |  |  |
| Taxon | Mean contribution | sd | Control | Disturbed | Cumulative contribution |
| Ulvoids | 0.14 | 0.15 | 8.62 | 28.49 | 0.30 |
| *Lithothamnion* sp. | 0.07 | 0.06 | 13.44 | 4.93 | 0.46 |
| *Gelidium* sp. | 0.04 | 0.04 | 7.60 | 4.07 | 0.55 |
| *Pyropia* spp. | 0.04 | 0.05 | 3.55 | 6.02 | 0.63 |
| *R. verrucosa* | 0.03 | 0.05 | 3.22 | 6.44 | 0.70 |
| *H. lecannellieri* | 0.02 | 0.03 | 5.65 | 3.51 | 0.76 |
| *C. officinalis* | 0.02 | 0.03 | 4.75 | 1.64 | 0.81 |
| *Polysiphonia* spp. | 0.02 | 0.03 | 3.58 | 2.44 | 0.85 |
| *Chaetomorpha firma* | 0.02 | 0.03 | 2.62 | 3.24 | 0.89 |
| *Verrucaria* sp. | 0.02 | 0.02 | 2.84 | 4.02 | 0.93 |
| **CHEU (*Perumytilus*)** |  |  |  |  |  |
| Taxon | Mean contribution | sd | Control | Disturbed | Cumulative contribution |
| Chthamalid barnacles | 0.28 | 0.20 | 54.97 | 3.47 | 0.49 |
| Ulvoids | 0.15 | 0.23 | 35.10 | 2.91 | 0.73 |
| *Pyropia* spp. | 0.07 | 0.11 | 9.69 | 9.75 | 0.86 |
| *Mazzaella laminarioides* | 0.06 | 0.10 | 6.44 | 9.82 | 0.96 |
| **CALF (*Perumytilus*)** |  |  |  |  |  |
| Taxon | Mean contribution | sd | Control | Disturbed | Cumulative contribution |
| Ulvoids | 0.23 | 0.28 | 59.00 | 6.34 | 0.39 |
| Chthamalid barnacles | 0.22 | 0.20 | 43.13 | 5.84 | 0.76 |
| *M. laminarioides* | 0.09 | 0.13 | 9.55 | 16.46 | 0.91 |
| **CHAI (barnacles)** |  |  |  |  |  |
| Taxon | Mean contribution | sd | Control | Disturbed | Cumulative contribution |
| Ulvoids | 0.22 | 0.21 | 2.21 | 5.28 | 0.45 |
| *M. laminarioides* | 0.09 | 0.10 | 2.31 | 3.95 | 0.62 |
| *R. verrucosa* | 0.06 | 0.10 | 8.73 | 6.85 | 0.74 |
| *Pyropia* spp. | 0.05 | 0.10 | 3.68 | 8.23 | 0.84 |
| *Nothogenia fastigiata* | 0.04 | 0.08 | 7.14 | 2.83 | 0.91 |
| **PUCA (*Perumytilus*)** |  |  |  |  |  |
| Taxon | Mean contribution | sd | Control | Disturbed | Cumulative contribution |
| Chthamalid barnacles | 0.24 | 0.21 | 2.40 | 38.64 | 0.43 |
| Ulvoids | 0.18 | 0.25 | 4.31 | 39.66 | 0.76 |
| *Pyropia* spp. | 0.06 | 0.10 | 5.85 | 8.14 | 0.86 |
| *R. verrucosa* | 0.05 | 0.11 | 1.67 | 9.56 | 0.96 |

Table A4. Pearson product-moment correlations (*r*) between functional and compositional forms of resistance, resilience, recovery, and invariability. Correlations were computed across sites and separately for each community type.

| Community | Resistance | | Resilience | | Recovery | | Invariability |
| --- | --- | --- | --- | --- | --- | --- | --- |
| *Mazzaella* | -0.54 |  | 0.53 |  | -0.27 |  | 0.07 |
| Barnacles | -0.10 |  | -0.52 | ***** | -0.57 | ***** | 0.10 |
| *Perumytilus* | -0.06 |  | 0.30 |  | 0.17 |  | 0.06 |
| ‘*’ *P* < 0.05 | | | | | | | |

Table A5 Pairwise Pearson product-moment correlations (*r*) between stability dimensions.

Correlations were computed across sites and community types.

|  | Resistance |  | Resilience |  | Recovery |
| --- | --- | --- | --- | --- | --- |
| *Functional domain* | |  |  |  |  |
| Resilience | -0.38 | ****** |  |  |  |
| Recovery | -0.15 |  | 0.01 |  |  |
| Invariability | 0.39 | ** | -0.36 | ****** | 0.10 |
| *Compositional domain* | |  |  |  |  |
| Resilience | -0.15 |  |  |  |  |
| Recovery | 0.06 |  | 0.02 |  |  |
| Invariability | -0.13 |  | -0.19 |  | -0.08 |
| ‘**’ *P* < 0.01 | | | | | |
